# Brain Electrical Bioactivity Informational Specifics in Relation to Behavioural and Cognitive Functions in Children with Learning Difficulties

**DOI:** 10.1101/2022.12.30.522305

**Authors:** Nelli Tolmacha, Konstantin Pudvosky, Janis Vandans, Andrey Bondarenko

## Abstract

The current paper performs a comprehensive assessment of electroencephalogram (EEG) parameters and features of cognitive functions of the brain in children of preschool (6-7) and primary school age (7-8) years with learning difficulties are considered. Spectral and coherent EEG analysis is conducted as well as particularities of gamified orientation in the maze. Spectral analysis of the EEG in children with learning difficulties revealed a speed decrease in the formation of almost all indicators of alpha activity and an increase in the power of slow-wave activity in the EEG, which was more pronounced in the left hemisphere of the brain. EEG coherence analysis revealed early differences in the spatial organization of the coherence function both in EEG rhythms and the cerebral hemispheres. A high level of motor activity in the maze, fragmented perception, unstable attention, low volume of operational memory, and poor optimization of skills in mastering a new space in children with learning difficulties correlated with indicators of age-related neurophysiological immaturity in terms of EEG.

## Introduction

In recent years, the number of students experiencing learning difficulties has been steadily growing in all countries, and these are children from all social strata of society [1,2]. Learning problems in different age groups (6-9 years old) were found in 15-40% of children, according to various sources [3,4]. EEG characteristics of preschool age are critical to understanding the causes of school learning difficulties in learning in different age groups, which reflect the significant changes that have occurred in the anatomy and functioning of the child’s brain by the beginning of learning because the fine regulation of intercentral relations and interhemispheric asymmetry of the brain marked already in primary school age [5,6,7]. The researchers emphasize that it is at this age that various forms of deviant behavior begin to form [8]; therefore, the attention of researchers is drawn to the preschool and primary school age.

The topicality of this problem is also due to the requirements of life, because in modern conditions of technological progress with an ever-increasing pace of life, it became necessary to assimilate vast amounts of information, leading to great psycho-emotional stress in both children and adults.

### Current State

It is known that by the beginning of schooling, neurophysiological mechanisms undergo significant progressive transformations that ensure the formation of cognitive activity. This, first of all, refers to the organization of the system of perception and attention, which allows us to consider this stage of development as extremely important for improving the information processes that form the basis of cognitive activity [9,10]. At the same time, great importance has always been attached to the ratio of involuntary and voluntary attention prevailing at this age since both of them bear features of immaturity at this age [11]. According to modern ideas about the age norm (normal development), the development of a child’s body, the age from 4 to 7 years, is considered the period of the first childhood and the period of 8-11 years - the second childhood [12]. In the educational system, these are preschool and primary school ages. In our study, we took two close age groups of 6-7 years and 7-8 years (the end of one and the beginning of the second period of childhood) because for both parents and the school, the question remains how to objectively evaluate and understand how each child responds to cognitive loads and at what age of 6,7 or 8 systematic schooling should begin, this is especially important if parents see behavioral disorders and feel that the child is not ready for learning, and he is sent from kindergarten to school.

The mentioned age periods of childhood are considered in neurophysiology as critical and sensitive for the development and improvement of both the mechanisms of adaptation in various situations and the features of the holistic perception of complex sensory information. A discrepancy between the requirements for a child at school and his individual functional capabilities at these stages of development can lead to a slowdown and even failure of adaptation, cause neurotic reactions or persistent stress, and cause harm [13,14].

And although almost all children usually go to school willingly, but after a short period of time, many of them no longer want to participate in the learning process, moving away from it through distractions. Some children get up and walk around the classroom, talking to a neighbor, or daydreaming, etc.. All these actions are united by one thing - the child does not perceive information, and does not master it, because it is too complex for the degree of development of his brain at this stage. This leads to the loss of fragments of knowledge and further complicates the assimilation of new knowledge. Many schools have already created new, more lightweight training programs, remedial classes, etc. The problem is that, more often than not, a child enters such a class and/or such a program after failing the regular class requirements. This leads to changes in the child’s self-esteem, which further raises the probability of a failure of adaptation, with consequent stress and moral trauma of the psyche, with all further consequences [15,16].

At the same time, there is little research present about the specifics of brain organization, conducting a comparison between the cognitive activity and behavior in children aged 6-8 years old belonging to the norm population and the ones with various developmental disorders.

The age of 6-8 years is socially significant because it is during this period that the basic skills necessary for school education are formed; therefore, timely detection of neurophysiological deviations or specific features of the cognitive functions of the brain will allow timely prevention of the identified difficulties of school adaptation.

### Research Goal

To assess the functional features of electrical activity and cognitive functions of the brain in children with learning difficulties of preschool age 6-7 years and primary school age 7-8 years. To identify the earliest functional EEG markers for possible prediction of the nature of impairments to the cognitive resource of the brain.

### Research methods

Two age groups of children were examined: -80 kindergarten children 6-7 years old and 80 first-grade students aged 7-8. Each age group consisted of two subgroups: subgroup - the age norm of 30 children each: 18 boys and 12 girls. and subgroups of children with learning difficulties (50 people): 34 boys and 16 girls.

For 6-7 year olds, the first subgroup of the age norm includes children who, according to teachers and psychologists, willingly and easily mastered the preparatory program in kindergarten.

At 7-8 years old: children of the subgroup of the norm, easily mastered the school curriculum of the first grade. The second subgroups include children with learning difficulties who are under the supervision of neurologists from 4-6 years of age. The neurologist’s assessment pointed to learning difficulties, impaired activity, and attention.

The work was carried out in compliance with the conditions of Articles 5, 6 and 7 of the Universal Declaration on Bioethics and Human Rights, with individual informed consent and parents’ initiative for examination. In accordance with the purpose of the work, examinations of children were carried out in the following order.

The brain’s electrical activity (EEG) was recorded on an electroencephalograph - EEG analyzer - 21/26, “Encephalan 131-03”, Medicom MTD, Russia. The electrodes were applied according to the “10-20.” system. A computer EEG was recorded in a monopolar ipsilateral ear lead in compliance with the standard conditions for this type of neurophysiological diagnostics in accordance with the following basic requirements: “standardization of external conditions and examination procedures, creation of an optimal psychological climate and a motivated set of subjects” [17]. For all children with learning difficulties and children of the age norm, EEG was recorded in conditions of quiet wakefulness with eyes closed at rest - up to 5 min. further functional loads, including - the solution of arithmetic problems-addition in mind. The average duration of the study was 15-20 minutes. Quantitative assessment of electroencephalograms and their compliance with the biological age norm was carried out based on the results of spectral and coherent analyzes, which were carried out using the built-in functions of mathematical data processing, the Medicom Mtd FAM software package. The method of calculating the frequency spectrum of the EEG power is one of the leading indicators of the process under study in clinical and scientific studies [19,20].

EEG coherence analysis, a relatively newer method, allows one to assess the degree of similarity of compared signals in a specific frequency spectrum between two leads or one with all leads [21]. The EEG was studied by epochs corresponding to the time intervals of the described procedures. The coherence dynamics made it possible to assess the relationship’s stability between the two processes’ corresponding frequency components [22].

The level of coefficient of coherence (CC) is a normalized value; its values lie within 0-1.00. When CC is less than 0.30, the coherence level is considered low. Values of 0.31-0.50 correspond to moderate; 0.51-0.70 is significant, and 0.71-1.0 is a high level of coherence (23.24). Our study considered significant and high coherence levels (reference interval 0.51-1.00), which made it possible to evaluate the features and degree of similarity of electrical wave processes recorded between the electrodes on the left and right hemispheres. The values of all types of coherence for each subject were determined in five frequency ranges: delta (0.5–4.0 Hz), theta (4.0–8.0 Hz), alpha (8.0–13.5 Hz), beta 1 - (14.0-22.0Hz), beta 2 - (24.00-35.00). Based on the results of the quantitative analysis, tables, histograms, and graphs were created. The data obtained were subjected to visual and mathematical analysis.

Indicators of children’s behavior (the cognitive resource of the brain) were assessed during the game - orientation in closed labyrinths of 2 degrees of complexity. Each movement in the maze was recorded and equated to one bit of incoming information in the form of nu-grams. Analysis of the results of the search/orientation activity was carried out according to Shannon 1987, the methodology modification is described in detail by Nikolskaya K. and Voronin L. 1996-98, and in 2005 and 2010, it has been updated by the authors. They assessed how each child understands the instruction (its arbitrary retelling), “how you need to navigate the labyrinth open on the computer screen to win a prize”. After the child visually memorized the path through the maze, only the open entrance to the maze remained on the screen, and the visual route through the maze was closed. It was necessary to move along the corridors of the labyrinth independently using the arrow keys, recalling the direction of movement from memory. Three visual cues were allowed (the labyrinth opened, and it was clear which section of the route to the goal the child was at the moment). The program made it possible to automatically register motor activity along the path and new movements and repeated ones, as well as the time spent in each corridor leading to the prize or the exit.

The nature of the child’s behavior in the maze was assessed: the level of search activity, the characteristics of perception and attention, the number of errors, the amount of memory, the characteristics of motor activity, its speed and purposefulness, the presence of skill optimization as the orientation in the maze, as well as the individual ability to form a search and acceptance algorithm: solutions, availability of a search strategy. The individual characteristics of the motivated and emotional components were recorded. Testing took 20 to 40 minutes, depending on the child’s computer skills. The latter had the most significant influence on the indicators of the speed of motor activity in the maze.

The results of all studies were statistically processed. Comparative assessment of quantitative characteristics was performed using parametric statistical criteria (Student’s t-test), and intergroup comparison of significance with the non-parametric distribution of related parameters was performed using the paired Wilcoxon test, and with disconnected samples, the Mann-Whitney test. The critical significance level of the tests was determined at p ≤ 0.050.

The results of the examination were compared with the biological age norm. The research center has a normative database on cognitive indicators of orientation in labyrinths in children in the norm from 6 to 15 years.

### Research Results

Quantitative data analysis on the relative values of the EEG power spectra in children of the two examined groups with learning difficulties revealed significantly higher values of the EEG power in the frequency ranges of slow waves compared to the age norm.

A comparison of the relative power values for all EEG rhythms in the two studied age groups of the norm and children with learning difficulties, respectively, is presented in Fig. 1 in the form of histograms and graphs.

**Figure. 1.**
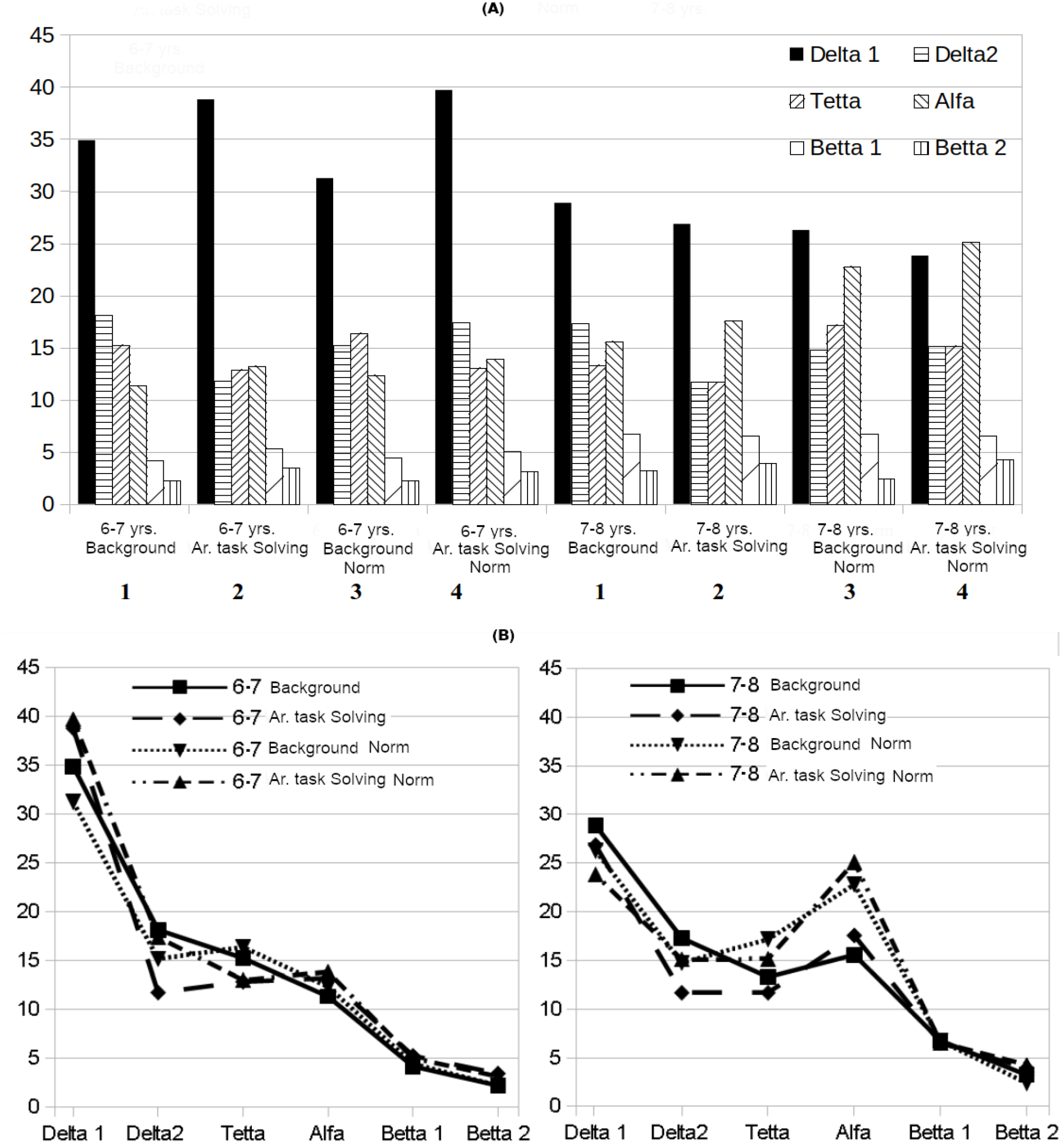
(A) - histograms of relative power values of all EEG rhythms in the ranges of delta 1,2, theta, alpha and beta 1,2 waves in children of two age groups 6-7 and 7-8 years old with learning difficulties -1,2 and their age norm -3,4; respectively, in the background state - 1,3 and during the test (addition in mind - arithmetic task solving) - 2,4 in each of the groups. (B) graphs * - significant age differences p<0.05; differences - at the trend level, p <0.1.

According to the data presented in Fig. 1, the following conclusions can be drawn: for children of two age groups with learning difficulties 6-7 and 7-8 years old, high EEG powers are characteristic in the ranges of slow rhythms delta 1-(0.50-2.00 Hz), delta 2-(2.00 -4.00 Hz) and theta - (4.00-8.00 Hz) waves both at rest and when solving an arithmetic problem (addition in mind) p<0.05. At the same time, significantly lower power values in the delta 1,2, ranges are recorded in 7-8 year old children of the norm than in 6-7 year old children of the norm. Comparison of the power of the EEG spectra in the ranges of slow waves in the cerebral hemispheres revealed a significant increase in the latter in the left hemisphere (p<0.05), which led to cortical asymmetry in delta activity from 30-65% (Fig. 1).

In children of both groups 6-7 and 7-8 years of age with learning difficulties, the power of alpha waves was significantly reduced both at rest-background and when solving arithmetic problems (p<0.05). At the same time, the slow formation of almost all indicators of alpha activity compared to the age norm is characteristic: low index, high amplitudes, irregularity of alpha waves, absence or randomness of modulation in amplitude, often a violation of the “spatial-zonal” distribution of alpha waves (p<0.05).

In 31% of 6-7 year old children with learning difficulties, alpha waves were recorded in the brain’s frontal regions. Interhemispheric asymmetry in amplitude for alpha activity ranged from 30-60%, and in 23% of children, a slight asymmetry was registered in frequency. The organization of alpha waves varied from moderately to significantly disorganized, and the modulation of alpha waves in amplitude in the EEG was absent in 73% of cases. In children, 7-8 years old with learning difficulties, the lack of formation of alpha activity indicators was less pronounced, but significant differences with the age norm remained. All EEG alpha activity indices in children of the two surveyed age groups with learning difficulties compared with their age norm corresponded to a younger age. In children of both groups 6-7 and 7-8 years of age with learning difficulties, the power of beta waves was significantly reduced both at rest-background and when solving arithmetic problems (p<0.05).

In children of the age norm at 6-7 years old, in the background EEG, a tendency to an increase in the power of beta 1 and beta 2 waves was revealed; and in 7-8 year old children, there is already a significant increase in these waves at rest, and when solving an arithmetic problem (p<0.05). At the same time, in children with learning difficulties, especially in 6-7 years old beta 1, beta 2, and gamma could be registered in the form of short discharges or were expressed locally in the occipital, temporal, or frontal regions of one of the hemispheres.

A coherent analysis of the EEG in all children of the studied age groups was performed according to indicators - the values of the coherence level (CC). A pronounced individual instability and variability were revealed. For the current study we have chosen average levels of EEG coherence across all frequencies (Fig. 2(A)), and coherent analysis across major interhemispheric longitudinal chains (right hemisphere: FP2-F4, F4-C4, C4-P4, P4-O2, FP2-F8, F8-T4, T4-T6, T6-O2; left hemisphere: FP-F3, F3-C3, C3-P3, P3-O1; FP1-F7, F7-T3, T3-T5, T5-O1) (Fig. 2(B)).

**Figure. 2.**
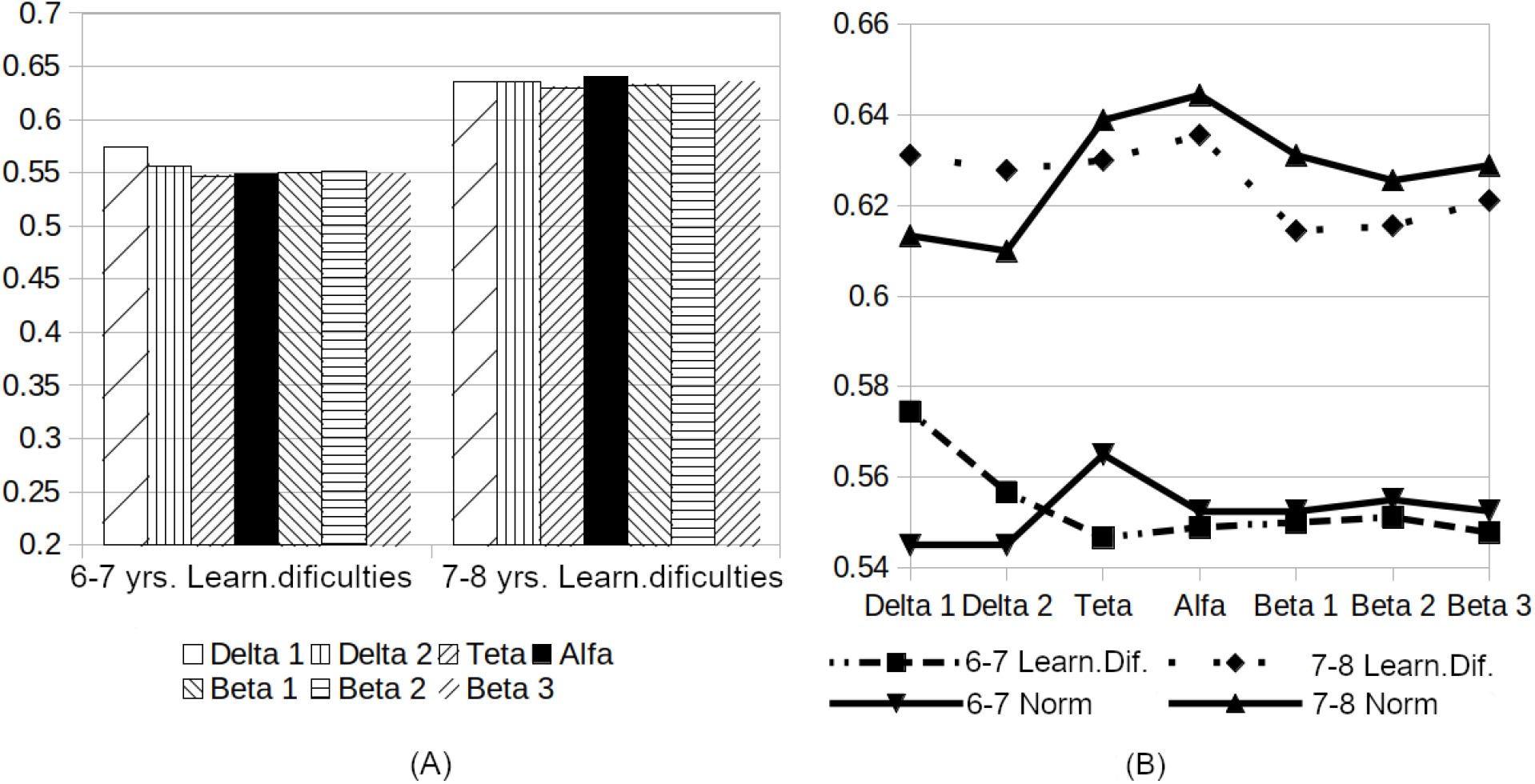
Average levels of the EEG coherence function within the reference corridor (0.51 -1.00) in children of two age groups 6-7 and 7-8 years old. The age norm groups are - 2,4; groups with learning difficulties -1,3; between vertically located leads in the left and right hemispheres of the brain (inside hemispheric), in the ranges of delta 1,2, theta, alpha, and beta 1,2,3 EEG waves. (A) - Average levels of coherence by EEG frequencies within groups having learning difficulties; (B) - The overall level of EEG spectra coherence according to standard inside hemispheric chains across all four groups of children.

According to the data presented in Fig. 2, we can draw the following conclusions: children with learning difficulties 6-7 years old in comparison to the same age group norm, compared with 7-8 year old children of both groups, pronounced differences are observed in the average level of the coherence function in all EEG wavebands (p <0.05). Thus, in 6-7-year-old children, the CC value was recorded in a narrow range: at rest, it varies from 0.51 to 0.58, and with a cognitive load from 0.55 to 0.65 - a significant level of coherence, then in 7-8 year old children, respectively, from 0.63 to 0.64 at rest and from 0.63 to 0.66 during exercise - a high level of coherence. At the same time, in children of both age groups with learning difficulties -1,3, redundancy of connections was revealed both at rest and on cognitive load, which in 6-7 year old children is expressed in the band of delta1,2 waves, and in 7-8 year old children in the bands delta2 and theta waves. These data on CC values correlated with the average values of the relative power of slow EEG rhythms. The level of coherence in the theta wave range, as well as the values of the relative power of theta waves in the background EEG in children of two age groups 6-7 and 7-8 years old with learning difficulties, were significantly lower than in the groups of their age norm (p < 0.05). It is characteristic that a significant level of the coherence function in the range of alpha and fast waves beta 1 and beta 2 is increased only in the normal groups; a trend was noted in 6-7 year old children, and a significant increase in cognitive load in 7-8 year old children (p<0, 05).

In groups of 6-7 and 7-8 year old children with learning difficulties compared to the age norm, a significantly higher variability of the coherence function of vertical connections in each of the cerebral hemispheres was registered both in the state of calm wakefulness and during the solution of an arithmetic problem. At the same time, a high level of coherence functions was registered in the band of slow waves (delta 1, delta 2, theta waves, in both hemispheres of the brain), mainly in the anterior-frontal-frontal and anterior-temporal-temporal areas of the cortex, and less often in the central-parietal-occipital areas. The coherence functions in the band of alpha and beta waves were the most variable and were found between closely spaced leads (short chains) such as Fp2-F8 and Fp1-F7, anterior frontal and anterior temporal leads of both hemispheres, as well as T4-T6; T6-O2 temporal and temporoccipital leads of the right hemisphere, and between the temporal T3-T5 and C3-P3 central parietal leads of the left hemisphere.

Connections between closely located points of the cortex are most pronounced in different frequency ranges in the right hemisphere, both at rest and when solving an arithmetic problem, and are more often recorded in the lateral leads of both hemispheres. In 6-7 year old children, short chains are recorded, in 7-8 year old children, especially in the age norm group, elongated chains of connections are recorded, for example, between Fp2-F8 -T4-T6; Fr1-F7-T3-T5 along the lateral leads, as well as between F4-C4-P4; C3-P3-O1 in the middle leads, the number of which significantly increases in the band alpha and especially beta 1 and 2 waves (p <0.05).

Consequently, in children of preschool (6-7) and primary school age (7-8) years old who are very close in age with difficulties in learning, characteristic age-related changes in both indicators, the power of electrical activity in various EEG rhythms, and early differences in the spatial organization of the function were registered. coherence in the cerebral hemispheres in different frequency ranges of the EEG.

An analysis of the behavior of both age groups of children with learning difficulties revealed that when they read the instructions about “playing” in the maze, 6-7-year-old children more often than 7-8-year-old children have problems understanding the instructions. However, considering the very strong desire of children to play, orientation in the maze after EEG registration was allowed for everyone. Three types of behavior were identified in children from 6 to 8 years old with learning difficulties.

The first type - all children of both age groups who had practically no computer skills could not retell the instruction except for the word “prize”. All attempts to teach the child how to move were impatiently rejected. Entering the labyrinth, their joyful, excited mood abruptly changed to a state of anxiety, but they still ignored the help, they began to move very slowly and uncertainly, and despite the encouragement, they themselves quit the game at the first stage, they could burst into tears.

The second type - the children had computer skills when learning what conditions, how to move, and what the purpose of the game - confidently answered: “everything is clear” - moved actively in the maze and already stubbornly refused contact, moved chaotically, making multiple stereotypical movements -forward-backward, or they made left turns all the time or repeatedly moved along the “ring”, making many repeated movements that did not lead to a result, but they themselves could not switch, and this did not work out for the elders, they were disappointed by the lack of results and fatigue, and only then they themselves refused to play, completely. Children with the first and second types of behavior were excluded from the survey.

The third type - all the children had computer skills, the children were more or less in contact with them, it was possible to agree on the rules of the game, they memorized and could partially retell the instructions themselves or with tips, or answered questions about how to play to get prizes, with a finger on the computer screen, they could show how they would move to the first and second prizes if they did not see the path. In general, four boys and two girls were excluded from the group of 6-7 year old children, and three boys and one girl were excluded from the group of 7-8 year old children. Children of the age norm had no problems with contact and understanding of the game’s rules. Separate informative indicators of physical activity of normal children and children with learning difficulties when orienting in the maze are shown in Fig. 3.

**Figure. 3.**
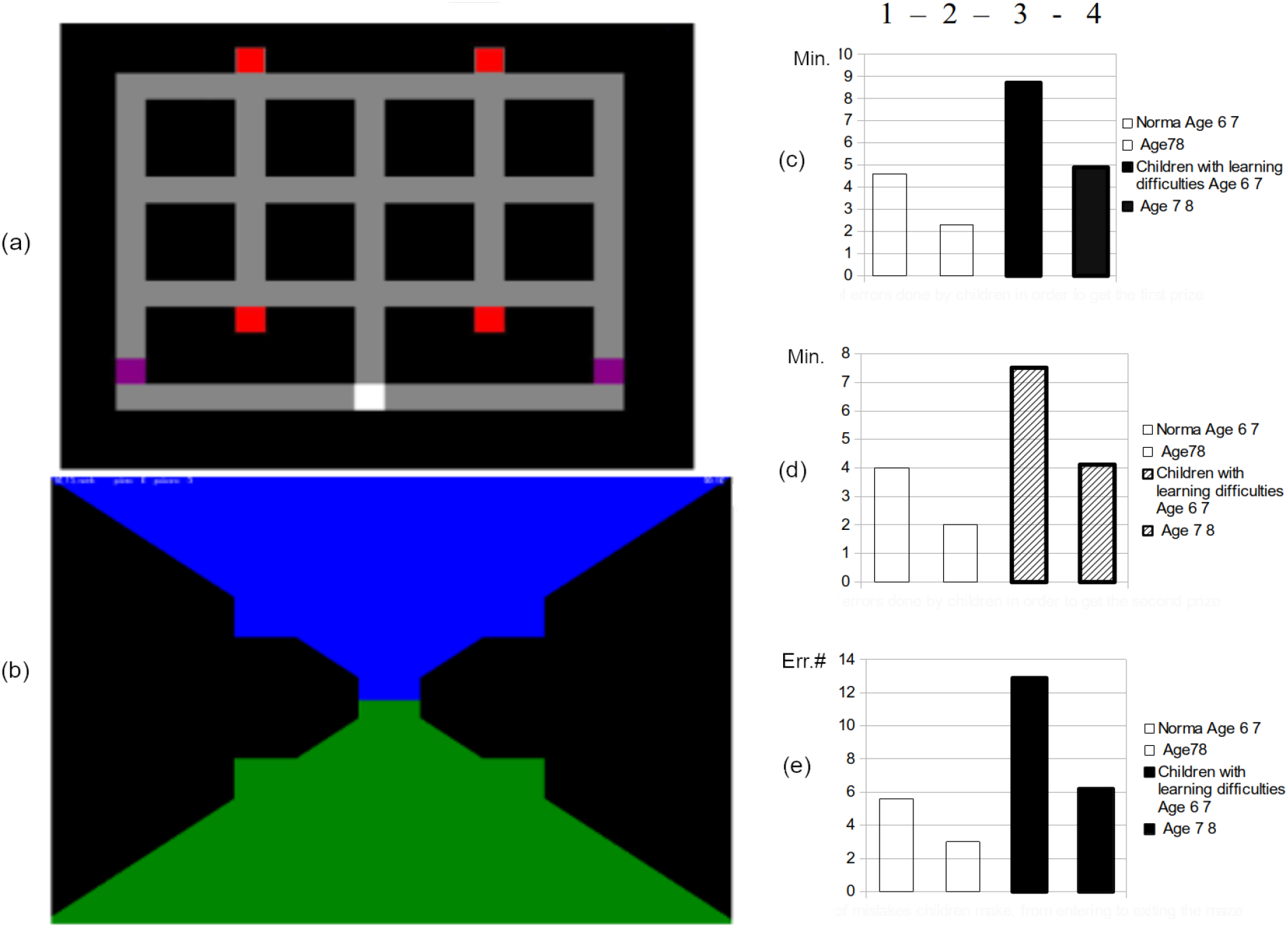
The results of motor activity in the maze in children of two norm groups: 1 - children 6-7 and 2 - 7-8 years old and two groups of children with learning difficulties: 3 and 4 respective age groups. (a) - maze plan, bottom red squares - traps, top red squares - prizes, white square - entrance, violet squares - exits; (b) - view of the labyrinth; (c) - the time spent (minutes) in the maze from maze entrance before receiving the first prize; (d) - the time spent (minutes) in the maze from first prize to the second prize and exit from the maze; (e) - the total number of errors (traps) while in the maze; (c,d,e) horizontal axis - groups. * - significant age differences, p<0.05; - differences at the trend level, p <0.1.

The results of orientation in the maze of children of two age groups of the norm showed that most children, especially 7-8 year olds, listen more carefully to the instructions and retell it in their own words when familiarizing themselves with the plan of the labyrinth, many say out loud the direction of movement “to the left”, “to the right”, some try to count the number of moves to the goal. At the beginning of the movement in the labyrinth, the keys that determine the direction of movement are more carefully evaluated.

In the first half of the survey, when orienting in the maze, children from the norm groups move more slowly, make fewer mistakes and unnecessary movements more carefully than children with learning difficulties; try to understand the meaning of their actions to achieve the result, especially after a hint, when the path they have traveled opens on the screen. Contact with an adult watching the game is normal for children in both groups; they are more ready to listen to the opinion of the elder and can stop the movement and switch their attention to the prompt.

For two groups of children with learning difficulties, a higher speed of motor activity is characteristic; they listened to prompts only at the very beginning of the game, and then, being carried away, they began to move chaotically, make mistakes, could make multiple stereotypical movements - forward-backward, repeatedly circling around, the movements become erratic. With great difficulty, especially 6-7 year old children were stopped and switched to view pictures on the computer screen and how they moved. The children calmed down very slowly and focused on viewing the path they had traveled, but it was only partially possible to analyze the features of the movement in 7-8 year old children. In the two groups, attention was maintained for a very short time, and only separate sections of the path of the labyrinth to the first prize were distinguished. As shown in Fig. 3(c), the time spent in the maze before receiving the first prize in children of both age groups of the norm (groups 1,2) is significantly less than in children with learning difficulties due to the large number of repeated and unnecessary movements that do not lead to the goal (groups 3,4) p<0.05. At the same time, if children from the norm groups in the second half of the game showed significantly less time to search for the second prize in the maze and a smaller number of errors p<0.05, and they were more independent in achieving a specific goal, then children with learning difficulties and in the second half of the game, they did a lot of repeated, superfluous, stereotyped movements that did not lead to a result (Fig. 3(e)). The development of a new space of the labyrinth, as well as the optimization of skills against the background of chaotic search activity, were sharply slowed down, and the total number of errors was significantly higher (groups 3,4) p<0.05 (Fig. 3(e)).

## Discussion

In the EEG of children 6-7 and 7-8 years old children with learning difficulties, compared with the age norm, the predominance of slow wave activity - delta1, 2 and theta in the frontal, central, parietal and occipital cortical leads along the internal chain of one, more often the left hemisphere or both hemispheres p<0.05. The two groups of children with learning difficulties were characterized by slow formation of almost all indicators of alpha activity: disorganization from moderate to significant, irregularity of waves, absence or randomness of modulation in amplitude, low index, and impaired zonal expression of alpha activity. The dynamics of the main EEG rhythms were due to age-related features of the maturation and functioning of brain structures [25]. The high level of power of slow waves and delayed maturation of the main signs of alpha activity in children with learning difficulties correlated with excessive formation of intra-hemispheric connections with a low level of coherence (up to 0.30) in the band of slow waves (delta-1.2, less often theta waves) in the frontal and temporal, less often in the central regions of the cortex of both hemispheres of the brain. Excess connections in the anterior areas of the cortex of both hemispheres of the brain were not found in children of two groups of the age norm. Previously, we recorded such a pattern of redundant connections in the anterior areas of the cerebral cortex in children with attention deficit hyperactivity disorder, a history of which perinatal disorders were described, most often associated with fetal hypoxia [26,27]. In both groups of children with learning difficulties, when interviewing parents, no perinatal disorders were noted (Apgar score - 8-9 points), according to the anamnesis, the children were under the supervision of neurologists.

Psychological examination of children with redundancy of connections of a low level of coherence revealed a high level of personal anxiety in both groups according to the method of Ch. Spielberger-Yu. An excessive number of connections in the anterior cortex in children slowed down the formation of significant connections and indicated the immaturity of the frontal and temporal cortex, due to individual characteristics of the maturation and functioning of brain structures [30]. Violation of the zonal power of the main EEG rhythms with an increase in the proportion of slow rhythms due to the immaturity of the functions of the frontal and temporal parts of the cerebral cortex in the early stages of ontogenesis were previously identified in children with minimal dysfunction [30], with aggressive behavior [31], with impaired reading and speech function [32], hearing impairment - hearing loss and in deaf children [33, 34]. The revealed structural and functional changes in children with aggressive behavior in the frontal and temporal lobes testified, according to the authors [35,36], to weak interhemispheric and intersegmental interactions of frontal cortex areas.

A comparative analysis of the identified EEG features in children of two age groups with learning difficulties compared with the age norm revealed differences in and the nature of their behavior. Lack of interest in the game, repeated repetition of the instruction, selectivity of its perception and unwillingness to contact and listen to the elders’ tips indicated that communication skills were not developed when communicating with the elders, especially among 6-7 year old children.

Orientation in the labyrinth revealed in these children unstable attention, perception of small segments of the path, low memory volume and poor optimization of skills in mastering the new space of the labyrinth, despite any emotional reinforcement. At the same time, a high rate of chaotic, stereotyped motor activity was recorded, which did not lead to a result, which indicated the predominance of processes of increased excitability in the nervous system and insufficiently formed control of voluntariness and purposefulness of movements. The nature of the behavior of children of two age groups with learning difficulties according to their ability in spatial orientation very clearly revealed the nature of their possible, future school difficulties in systematic learning, and the identified features of the EEG, apparently, were part of the main causes of these difficulties. The EEG pattern in these children in both age groups corresponded to younger age.

In terms of determining the child’s readiness for systematic learning at school, it is of interest to compare in more detail the EEG indicators and the characteristics of behavioral reactions when orienting in the maze between two groups of the age norm - preschool - 6-7 and primary school age - 7-8 years.

In the EEG of children of the age norm, against the background of a quantitative decrease in the severity of the power of slow waves in 7-8 year old children, compared with 6-7 year old children, a higher level of maturation of alpha activity indicators was revealed. Alpha waves are no longer sufficiently organized or organized, regular, the zonal-spatial distribution of alpha waves is formed, and their amplitude modulations are well expressed. EEG frequency ranges revealed early differences in the spatial organization of intrahemispheric connections and a higher level of their coherence across the cerebral hemispheres. In 7-8 year old children, compared with 6-7 year old children, elongation of short chains (Fp1-F7-T3; and Fp2-F6-T4-T6) was registered, their number is increased not only in the side chains, both in the left and right hemispheres, but also in both middle chains (F3-?3-P3-O1; and (F4-C4-P4-O2) of the cerebral cortex in 7-8 year old children compared to 6-7 year old children - a significantly larger number of single connections was also registered in hemispheres between closely located points of the cortex in different areas and in different frequency ranges, p<0.05. At the same time, while in 6-7 year old children CC values were recorded in a narrow range: significant differences both at rest and under cognitive load, then in 7-8 year old children CC was significant at rest, and a high level of coherence was already registered during exercise, which in the range of alpha waves correlated with an increase in the relative power of alpha activity, especially on cognitive load (p <0.05). A change in the nature of the spatial organization of the intrahemispheric connections of the brain and a significant increase in the level of coherence to a high level in the range of alpha and fast beta 1 and beta 2 waves on cognitive load in 7-8 year old children indicated an increase in functional connections between different parts of the cortex within each hemisphere, and consequently about the expansion of the functionality of the frontal, temporal, central and parietal symmetrical areas of the cortex (p<0.05).

The increase in the level of the coherence function in the range of fast waves beta 1 and beta 2 in younger schoolchildren at rest, and on cognitive load, according to Machinskaya R., reflects the level of activation of various areas of the cerebral cortex during the solution of cognitive tasks [37]. Taking into account the peculiarities of the behavior of 7-8 year old children in the maze, it should be noted that a significant increase and a high level of coherence, in the alpha wave range, not only at rest but also on cognitive load, which apparently can be assessed as one of the early signs of emerging arbitrariness behavior.

Consequently, in very close-aged children belonging to the norm groups at the ages of 6-7 years and 7-8 years, quantitative and qualitative changes in the electrical activity power indicators for various EEG rhythms are detected and early differences in the nature of the spatial organization of the EEG coherence function are revealed. The change in neurophysiological parameters correlated with the dynamics of the cognitive functions of the brain and the nature of behavior during the development of a motor skill in the maze.

Evaluation of the features of the studied EEG parameters and cognitive functions of the brain in children with learning difficulties of both preschool and primary school age revealed a slow, uneven functional maturation of the main age-related EEG parameters, the dynamics of which correlated with slowly and difficultly developed motor skills of orientation when mastering a new space in the labyrinth. The revealed lack of formation of communication skills, especially in 6-7 year old children, selectivity of perception and attention, low memory volume, and the nature of motor activity indicated a high level of activation of behavioral reactions, the predominance of excitation processes in the nervous system and low activity of inhibition processes.

Features of the formation and functioning of the nervous system in children with learning difficulties show that they are not ready for the mental loads of systematic learning.

## Conclusions

1. Comparative analysis of EEG parameters in children of children with learning difficulties of both preschool and primary school age - 6-8 years, and in children of the norm of the same age, respectively, revealed the following differences: Violation of the zonal power of the main EEG rhythms with an increase in the proportion of slow rhythms in the cortex of one or both hemispheres of the brain. Low level of coherence across all EEG rhythms. Slow formation of all indicators of alpha activity. A high level of variability of all rhythms due to the immaturity of the functions of the frontal and temporal parts of the cerebral cortex at this stage of ontogenesis.
2. The behavior of children with learning difficulties 6-7 and 7-8 years old in the maze was characterized by a low level of communication skills, high speed and difficult to switch between search and motor activity in the first and second half of the game, a large number of mistakes, repeated and unnecessary, stereotypical movements that did not lead to a result, which indicated the predominance of processes of increased excitability in the nervous system and insufficiently formed control, arbitrariness, and purposefulness of motor activity.
3. The basic functions of cognitive indicators of the brain in children with learning difficulties, 6-7 and 7-8 years old: unstable attention, fragmented selective perception, low memory volume, and poor optimization of goal search, which correlated with indicators of age-related neurophysiological immaturity according to EEG indicators, caused a low the level of formation of a motor skill in the maze, especially in 6-7 year old children, and testified to the unpreparedness of children for the loads of systematic learning.
4. The results of the functional analysis of EEG indicators are informative and promising in terms of assessing the maturation of neurons functional activity levels in all areas of the cerebral cortex in children in ontogeny in normal conditions and with various learning difficulties and can also be used as the earliest functional markers of the electrical activity of the brain to predict the nature of possible cognitive resource impairments or the behavior of the child in general.
5. EEG indicators that correlate with the characteristics of behavioral reactions and cognitive functions of the brain during orientation in the maze can be used as basic functional criteria for brain maturation in ontogenesis for the development of computer programs - express diagnostics of the degree of readiness of children for the systematic learning and timely prevention of possible school problems.

## References

1. Semyonova O.A., Machinskaya. Age-Related Transformations of Cognitive Functions in Cildren Aged 5-7 Years: A Neuropsychological Analysis. Культурно-историческая психология 2012, 2, pp.20–28.

2. Фесенко Е.В. Фесенко Ю.А. Синдром дефицита внимания и гиперактивности у детей-СПб: Наука и Техника 2010 -384c.

3. Кожушко Н.Ю., Евдокимов С.А., Матвеев Ю.К., Терещенко Е.П., Кропотов Ю.Д. Исследование Локальных Особенностей ЭЭГ у детей с нарушениями психического развития методами независимых компонент, Физиология человека. 2014. T. 40. № 5. C. 30-37.5.

4. Королёва Н. В. Становление биоэлектрической активности мозга у детей-дошкольников с факторами риска перинатальной патологии центральной нервной системы: Дисс. … канд. биол. наук. Иркутск, 2000.

5. Королева Н.В. Формирование биоэлектрической активности головного мозга у детей в онтогенезе / Н.В. Королева, С.И. Колесников. Иркутск: Издво Ирк. гос. ун-та, 2005. 88 c.

6. Скороходова Т.А. Функциональная организация интегративной деятельности мозга у детей младшего школьного возраста с разным уровнем интеллектуального развития: автореф. дис. канд. биол. наук. — T., 2001. — 16c.

7. Пономарева, Т. В Становление функциональных асимметрий в раннем онтогенезе автореферат диссертации. 2010. 165c.

8. Семенович А.В. Нейропсихологическая коррекция в детском возрасте. Метод замещающего онтогенеза.-М, Генезис, 2007 — 474c.

9. Machinskaya R.I. Neurophyziological mechanizms of volluntary attention: A review. Journal of Higher Nervous Aktiviti 2003.Vol.53. №2. 133–150.

10. Мачинская Р. И., Семенова О.А. Особенности формирования высших психических функций у младших школьников с различной степенью зрелости регуляторных систем мозга./ Ж. Эволюционной биохимии и физиологии. 2004.T. 40. №5

11. Kichigina V. F. Changes in Theta and Gamma Network Oscillations during the Development of Neurodegenerative Disorders (Review) Modern Technologijas Medicine /Vol. 11, Issue 1 (2019)→7

12. Pankov M. N., Kozhevnikova I. S., Sidorova E. Yu., Gribanov A. V., Startseva L. F. Psychophysiological Characteristics of Children with Aggressive Behavior. Ekologiya cheloveka [Human Ecology]. 2018, 2, pp. 37–44.

13. Sidorova E. Yu., Antonova I. V., Podoplekin A. N., Pankov M. N., Gribanov A.V. Neuroenergetics in children of primary school age with aggressive behavior. Ekologiya cheloveka [Human Ecology]. 2015, 2, pp. 51–56.

14. B. J. Casey, J. N. Epstein, J. Buhle et al., “Frontostriatal connectivity and its role in cognitive control in parent-child dyads with ADHD,” The American Journal of Psychiatry, vol. 164, no. 11, pp. 1729–1736, 2007.

15. A. Cubillo, R. Halari, C. Ecker, V. Giampietro, E. Taylor, and K. Rubia, “Reduced activation and inter-regional functional connectivity of fronto-striatal networks in adults with childhood Attention-Deficit Hyperactivity Disorder (ADHD) and persisting symptoms during tasks of motor inhibition and cognitive switching,” Journal of Psychiatric Research, vol. 44, no. 10, pp. 629–639, 2010.

16. Дубровинская Н.В., Фарбер Д.А., Безруких М.М. Психофизиология ребенка: Психофизиологические основы детской валеологии М.: Гуманит. изд. центр ВЛАДОС, 2000. 144 c. Дополн.2011.

17. Зенков Л.Р. Клиническая электроэнцефалография (с элементами эпилептологии) - М, МЕДпресс-информ, 2002 — 369c

18. Иванов Л.Б. Прикладная компьютерная электроэнцефалография — М. Научно-медицинская фирма МБН. 2000. 251c.

19. Гнездицкий В.В. Обратная задача ЗЗГ и клиническая электроэнцефалография (картирование и локализация источниковэлектрической активности мозга-М, МЕДпресс-информ, 2004. 624c.

20. Петрова Е.В. Пироженко. Спектральный анализ электроэнцефалограммы В кн Прикладная компьютерная электроэнцефалография 2000. 30–76c.

21. Кулаичев А. П. Об информативности когерентного анализа /А. П. Кулаичев // Журнал высшей нервной деятельности.- 2009. – T. 59. № 6. - C. 766–775.

22. Кулаичев А.П. Компьютерная электрофизиология и функциональная диагностика. 2016. M.- Форум-Инфра-M, 44C.

23. Рожкова Л. А. Межполушарная когерентность ЭЭГ детей младшего школьного возраста с нарушениями перинатального развития / Рожкова Л. А. // Актуальные вопросы функциональной межполушарной асимметрии и нейропластичности / под ред. С. Н. Иллариошкина, В. И.Кобрина, В. Ф. Фокина. М. : Научный мир, 2009. C. 418–423.

24. Яковенко Е.А., Чутко Л.С., Пономарев В.А., Сурушкина С.Ю., Никишена И.С., Кропотов Ю.Д. Особенности спектров мощности основных ритмов ЭЭГ у детей с разными типами синдрома дефицита внимания с гиперактивностью, Физиология человека. 2013. T. 39. № 1. C. 30.

25. Романчук Олег Синдром дефицита внимания и гиперактивности у детей.-М.Генезис, 2010 3.36c.

26. Толмача Н., Пудовскис К., Bанданс Я. Исследование электрической активности и когнитивных особенностей мозга для создания программы определения готовности ребенка к школе. Первый международный конгресс по клинической нейрофизиологии стран СНГ, ШОС,Азии и Европы. Алматы, Казахстан 3-6 октября 2018. -41–42c.

27. Pudovskis K., Tolmacha N. Functional maturity jf brain and readiness of children for systematic learning. 15 International Interdisciplinary Congress Neuroscience for Medicine and Psychology, 2015, 344–346.

28. Tolmacha N., Pudovskis K., Vandans J. Peculiarities of brain maturation for children with attention deficiency syndrome with hyperactivity. Fourth International Interdisciplinary Congress Neuroscience for Medicine and Psychology 2008, 396–398s.

29. Tolmacha N., Solomenikova I., Kazanovska I., Pudovskis K., Vandans J. Stress reactions peculiarities in pre-school and early school age children. Third International Interdisciplinary Congress Neuroscience for Medicine and Psychology. Ukraine, 2007, 234–235.

30. Ponomarev V.A., Grin-Yatsenko V.A., Kropotov J.D., Mueller A., Candrian G. GROUP INDEPENDENT COMPONENT ANALYSIS (GICA) AND CURRENT SOURCE DENSITY (CSD) IN THE STUDY OF EEG IN ADHD ADULTS // Clinical Neurophysiology. 2014. T. 125. № 1. C. 83–97.

31. Svyatogor I.A., Guseva N.L., Sofronov G.A., Sirbiladze K.T. Assessment of background and reactive EEG patterns in children with minimal brain dysfunctions // Medicinskij akademicheskij zhurnal SZO RAMN, 2013, tom 13, № 2, S.51–58

32. Smirnova T. P. Psikhologicheskaya korrektsiya agressivnogo povedeniya detei Psychological correction of aggressive behavior in children. Rostov-on-Don, Feniks Publ., 2007, 154 p.

33. Turner I. E. Gendernyi podkhod v profilaktike agressivnogo povedeniya mladshikh shkol’nikov (kand. dis.) [Gender-sensitive approach in the prevention of aggressive behavior of younger schoolboys (Cand. Diss.)]. Moscow, 2009, 155 p.

34. Подоплёкин А. Н., Сидорова Е. Ю., Антонова И. В. Распределение уровня постоянного потенциала головного мозга у детей 7—11 лет с высоким уровнем агрессивности // Журнал медико-биологических исследований. 2015. № 1. C. 49—57.)

35. Косарева Екатерина Игоревна. Исследование достоверных отличий параметров фоновой ЭЭГ у детей с нарушениями слуха // Наука через призму времени.-2018.-№12 (21). Кулаичев, А. П. Об информативности когерентного анализа /А. П. Кулаичев Журнал высшей нервной деятельности.- 2009. – T. 59. № 6. - C. 766–775.

36. Efficient Of Inter-Hemispheric Coherence As The Indicator Of Brain Integrative Processes At Healthy People Of Youthful Age, 2014 N.A. Shnayder, V.G. Nikolaev, N.N. Medvedeva, E.E. Vasilyeva, O.B. Pen2, K.A. Gazenkampf, Yu.B. Govorina, A.A. Romanenko, S.N. Derevtsova Proceedings of the Samara Scientific Center of the Russian Academy of Sciences, volume 16, No. 5 (4), 2014

37. Changes in frontal EEG coherence across infancy predict cognitive abilities at age 3: The mediating role of attentional control. Margaret Whedon, Nicole B Perry, Susan D Calkins, Martha Ann Bell 2016 Sep;52(9):1341–52. doi: 10.1037/dev0000149. Epub 2016 Jul 21.

